# Combined Selective Plane Illumination Microscopy and FRAP maps intranuclear diffusion of NLS-GFP

**DOI:** 10.1101/2020.05.25.114827

**Authors:** Chad M. Hobson, E. Timothy O’Brien, Michael R. Falvo, Richard Superfine

## Abstract

Since its initial development in 1976, fluorescence recovery after photobleaching (FRAP) has been one of the most popular tools for studying diffusion and protein dynamics in living cells. Its popularity is derived from the widespread availability of confocal microscopes and the relative ease of the experiment and analysis. FRAP, however, is limited in its ability to resolve spatial heterogeneity. Here, we combine selective plane illumination microscopy (SPIM) and FRAP to create SPIM-FRAP, wherein we use a sheet of light to bleach a 2D plane and subsequently image the recovery of the same image plane. This provides simultaneous quantification of diffusion or protein recovery for every pixel in a given 2D slice, thus moving FRAP measurements beyond these previous limitations. We demonstrate this technique by mapping intranuclear diffusion of NLS-GFP in live MDA-MB-231 cells; SPIM-FRAP proves to be an order of magnitude faster than fluorescence correlation spectroscopy (FCS) based techniques for such measurements. We observe large length-scale (> ~500 nm) heterogeneity in the recovery times of NLS-GFP, which is validated against simulated data sets. 2D maps of recovery times were correlated with fluorescence images of H2B to address conflicting literature on the role of chromatin in diffusion of small molecules. We observed no correlation between histone density and diffusion. We developed a diffusion simulation for our SPIM-FRAP experiments to compare across techniques; our measured diffusion coefficients are on the order of previously reported results, thus validating the quantitative accuracy of SPIM-FRAP relative to well-established methods. With the recent rise of accessibility of SPIM systems, SPIM-FRAP is set to provide a simple and quick means of quantifying the spatial distribution of protein recovery or diffusion in living cells.

**Statement of Significance:** We developed selective plane illumination microscopy combined with fluorescence recovery after photobleaching (SPIM-FRAP) to perform simultaneous FRAP measurements for each pixel in a 2D slice. This technique has the potential to be implemented on almost any light sheet microscope with minimal software development. FRAP studies were previously unable to resolve spatial heterogeneity and FCS techniques require minute-long acquisition times; SPIM-FRAP remedies both of these issues by generating FRAP-based diffusion maps in 4 seconds. This technique can easily be expanded to 3D by photobleaching a single plane and performing light sheet volumetric imaging, which has the benefits of minimal photobleaching and phototoxicity for studying long-term protein turnover. Furthermore, SPIM-FRAP of slowly-recovering structures enables characterization of spatial distortions to measure intracellular stresses.

## Introduction

Fluorescence Recovery After Photobleaching (FRAP) (1) is one of the most prevalent techniques for studying intracellular diffusion and protein dynamics. In brief, a region of interest of a fluorescently labeled sample is exposed to a high-intensity light source, thus photobleaching this specific region. Either through diffusion or (un)binding, the fluorescence of the bleached region recovers, allowing one to understand both the timescales of the recovery as well as the (im)mobile fraction. The widespread availability of both fluorescent proteins and point-scanning confocal microscopes has dramatically increased the accessibility of performing FRAP experiments. The other most common technique for studying such dynamics is fluorescence correlation spectroscopy (FCS) (2). FCS takes advantage of the fluorescence intensity fluctuations resulting from diffusion in and out of an excitation volume, and uses correlation analysis to extract quantitative measures of diffusion (i.e. the diffusion coefficient). FRAP and FCS have dramatically accelerated research in the realm of diffusions and protein dynamics.

The relative merits of FCS and FRAP are well documented. Both FRAP and FCS require precise knowledge of the illumination volume for an accurate measurement of the diffusion coefficient, often making absolute quantification difficult (3). Although, descriptions of relative changes of intracellular dynamics under various interventions are still readily possible. Beyond diffusion measurements, FRAP can measure immobile fractions, while FCS can measure absolute concentration. As for limitations, FRAP can be sensitive to bleach correction, while FCS requires careful consideration of the concentration of the fluorescent protein of interest and has higher signal-to-noise requirements than FRAP (4). However, the primary drawback of FRAP is the inability to distinguish heterogeneity of diffusion in a given sample. That is, each FRAP experiment produces one measurement for a given region of interest. Investigators have looked to FCS for studying such heterogeneous dynamics. By either iteratively performing FCS measurements across a cell of interest or using a selective plane illumination microscopy (SPIM) system coupled with FCS (SPIM-FCS), investigators have been able to generate 2D maps of intracellular diffusion (5–11). These measurements require acquisition times on the order of minutes to create such maps, presumably due to the high signal-to-noise requirements of FCS. Here, we address these limitations of both FRAP and FCS by combining SPIM with FRAP (SPIM-FRAP) to generate simultaneous 2D maps of intranuclear diffusion.

Previous work has used a SPIM microscope to image after photobleaching with a focused beam (12). Here, we use the light sheet to both photobleach and image our sample. By photobleaching a single plane that coincides with our image plane, we still allow for diffusion into the image plane from the rest of the 3D volume. Each pixel in our image then provides a simultaneous FRAP measurement. We demonstrate our SPIM-FRAP technique by mapping the diffusion of NLS-GFP in MDA-MB-231 cells with improved temporal resolution over FCS-based techniques (SPIM-FRAP: 4 seconds, FCS-based: ~minutes) and the added spatial information compared to traditional FRAP. This decrease in acquisition time in not a feature of the specific experiment at hand, but rather is indicative of fundamental limitations of FCS and FRAP. For accurate FCS measurements, the acquisition time should be no less than 100*τ*_*D*_, where *τ*_*D*_ is the relevant time scale given by *τ*_*D*_ = *A*_*eff*_/4*D*, *A*_*eff*_ is the effective area, and *D* is the diffusion coefficient (13). Accurate FRAP measurements require only acquisition times of ~10*τ*_*D*_. the observed order of magnitude gain in acquisition time of SPIM-FRAP is then a fundamental feature of the technique itself. We further simulate SPIM-FRAP experiments to determine what degree of heterogeneity can be detected by our technique. To convert recovery times to diffusion coefficients, we generate a simulation of diffusion that accounts for diffusion during the bleach pulse. Both the diffusion coefficients we report are consistent with previous FRAP and FCS literature.

## Results and Discussion

SPIM-FRAP experiments were performed by photobleaching a single plane with a custom light sheet microscope (Fig. 1A) of live MDA-MB-231 cells co-expressing either NLS-GFP and H2B-mCherry or NLS-GFP and 53BP1-mCherry. We then use the same light sheet at a reduced power to image the photobleached plane as the NLS-GFP intensity recovers via diffusion into the image plane (Fig. 1B, Movie S1). Images were bleach corrected before analysis. The recovery images were minimally blurred in FIJI (14) with a single-pixel Gaussian blur prior to analysis. An exponential recovery of the form

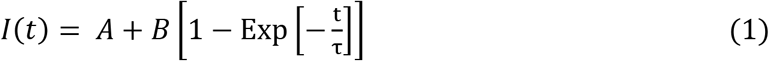

was fit to the intensity of each pixel during the recovery portion of the time series (Fig. 1D). From these data we can extract a characteristic recovery time, τ, for every pixel in the image plane. This subsequently provides a 2D map of τ for intranuclear diffusion of NLS-GFP (Fig. 1C). The total acquisition time for each experiment was 4 seconds, providing an order-of-magnitude improvement to FCS-based techniques (5–11).

**Figure 1.**
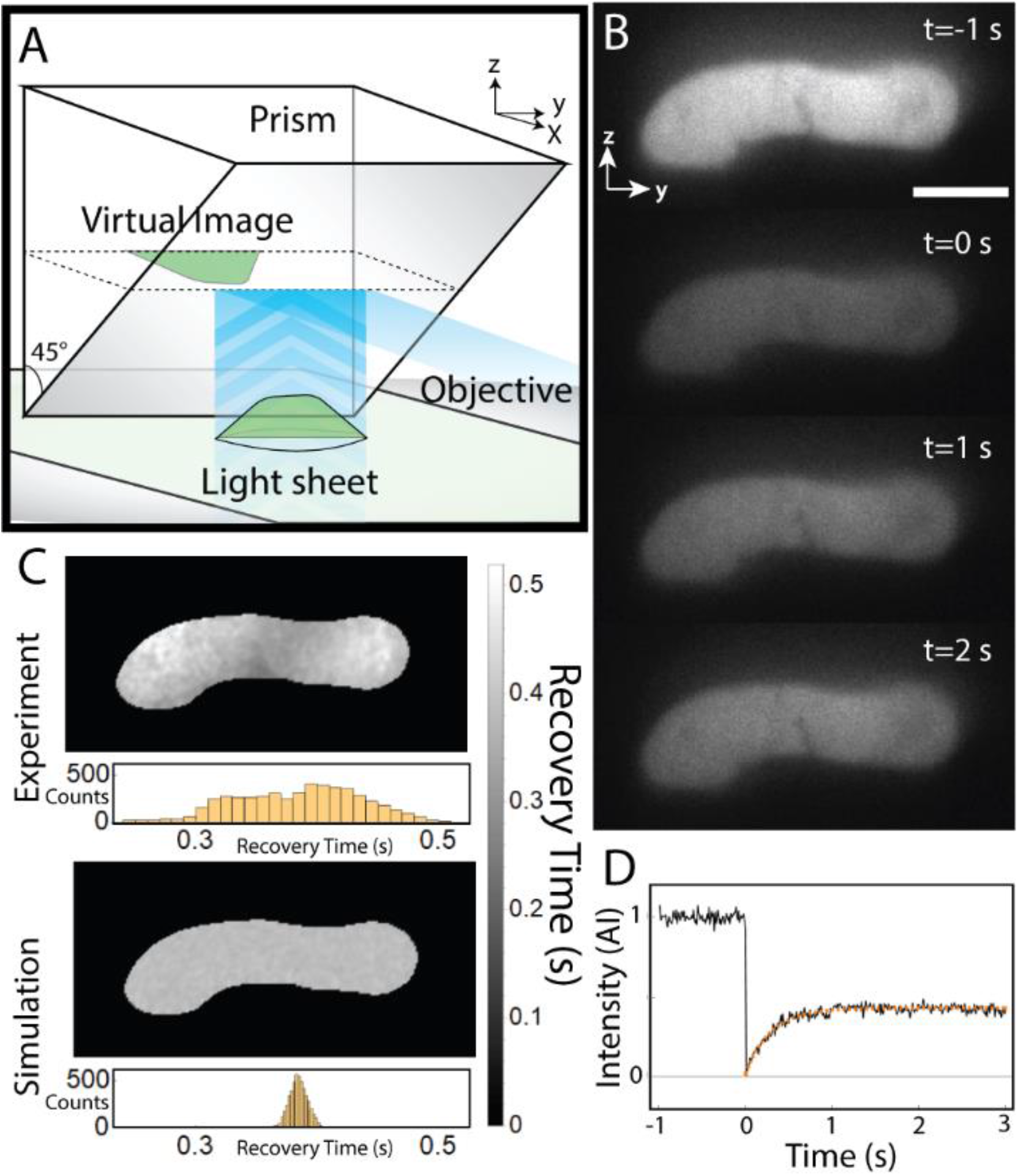
SPIM-FRAP generates simultaneous 2D maps of intranuclear NLS-GFP recovery times. (A) Schematic of our custom, single-objective SPIM microscope. (B) A side-view SPIM-FRAP image sequence of NLS-GFP showing the simultaneous recovery across the entire image plane (Movie S1). Scale bar = 5 μm. (C) 2D maps and histograms of NLS-GFP recovery times for a SPIM-FRAP experiment and a simulated data set. The ground truth of the simulation is a uniform recovery time. Noise and Gaussian blurring prior to curve fitting are accounted for in the simulation. (D) Time series of a single-pixel intensity of the image sequence shown in (B) after Gaussian blurring. The recovery is bleach corrected and the orange curve represents an exponential recovery fit.

Before drawing conclusions regarding the spatial distribution of *τ*, we sought to determine what our method is able to resolve. To study this, we generated a suite of simulated SPIM-FRAP data sets as described in the Supplementary Material (Fig. 1C, S1). Simulations with a uniform recovery time showed that the addition of noise to the recovery curve broadened the distribution of measured recovery times, and that our 1-pixel Gaussian blur served to tighten this distribution at the cost of spatial resolution (Fig. 1C). Extracting *τ* for every pixel, then, does not necessarily mean that we have single-pixel resolution for discerning structure in recovery maps. Additionally, we simulated recovery with 5×5 (540×540 nm) pixel squares that were prescribed a 20% faster recovery time than the rest of the data set. Our analysis was still able to easily detect this structure. We then concluded that the short length scale (~2-3 pixels, ~216 – 324 nm) mottling structure is an artifact of the Gaussian blur coupled with noise in the image sequence. However, the large length-scale (> 5 pixels, 540 nm) heterogeneity in the experimental recovery time map is not an artifact of the analysis. This implies there is a spatial dependency of intranuclear diffusion of NLS-GFP, consistent with FCS measurements (8). This distribution was, in some instances, bimodal. The source of this heterogeneity is presently unknown and warrants further studies. Intranuclear phase separated nucleoli have been shown to be more viscous than the surrounding nucleoplasm (5). The observed heterogeneity in diffusion could then be indicative of liquid-liquid phase separation in the nucleus (15). Alternatively, these data could suggest that the heterogeneity of the viscosity of the nucleoplasm is due to variations in concentration of macromolecules. This could have profound effects on nuclear mechanical properties and mechanotransduction (16). Finally, binding of the NLS to RNA could be a source of varied diffusion (17). Regardless of the origin, the observed spatial heterogeneity of intranuclear diffusion highlights that intranuclear transport is similarly heterogeneous on >~500 nm length scales. To understand the potential origin of this heterogeneity, we sought to correlate our diffusion measurements with nuclear structure.

Previous FCS literature has suggested that the diffusion of small molecules through the nucleus has little correlation with chromatin concentration (8); further work, however, reports that through use of an FCS variant using pair Correlation Functions (pCF) that DNA does indeed play a role in hindering transport of small molecules (18). SPIM-FRAP provides an opportunity to address such questions as it allows for correlation of recovery time maps with fluorescence images of other structures. By collecting a fluorescence image of H2B-mCherry prior to performing a SPIM-FRAP experiment of NLS-GFP on the same image plane (Fig. 2A), we can explore any correlation between histone density and diffusion. For each cell examined (n=11), we plotted the normalized H2B intensity versus the measured recovery times per pixel (Fig. 2B) and calculated the correlation coefficient (Fig. 2C). We observed a large spread in the correlation coefficients with no significant difference from zero correlation. This implies, similar to previous work (8), that there is no immediate spatial correlation between histone density and intranuclear diffusion of NLS-GFP. However, this does not negate some of the more intricate theories regarding barriers to long-range diffusion and sudden bursts of motion across dense regions of DNA (18). Additionally, SPIM-FRAP may not be able to detect a correlation on a length scale of ~100 nm. We further treated MDA-MB-231 cells co-expressing NLS-GFP and 53BP1-mCherry with Trichostatin A (TSA) to decondense interphase chromatin levels before performing SPIM-FRAP experiments. The peak of each recovery time distribution was determined by fitting a Gaussian curve to the respective histogram; for bimodal distributions, two peak recovery times were extracted. We observed no significant difference in peak recovery times for WT and TSA-treated cells (Fig. 2D).

**Figure 2.**
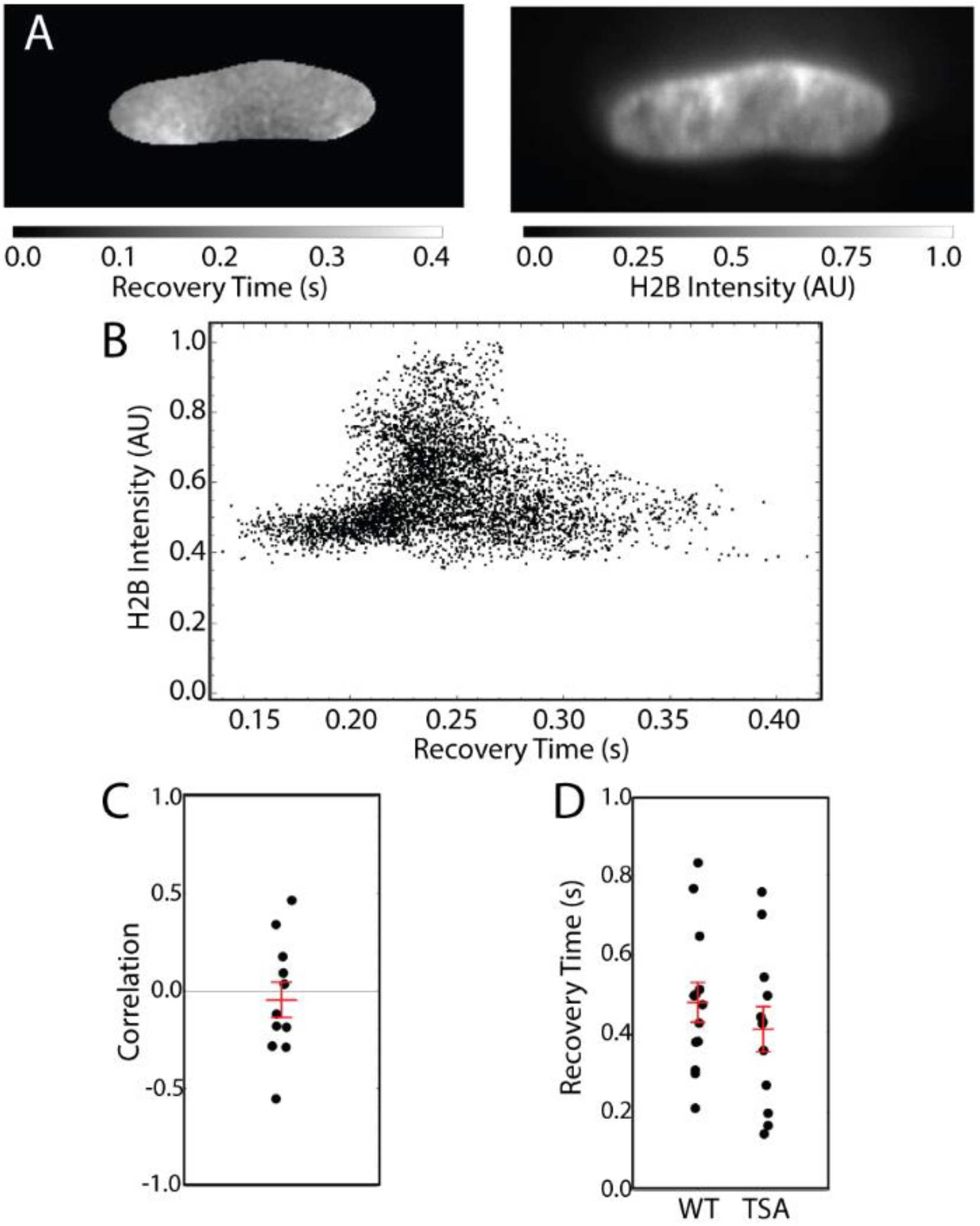
Chromatin Density does not inhibit diffusion of NLS-GFP. (A) SPIM-FRAP recovery time map for NLS-GFP and a corresponding image of H2B-mCherry. (B) A plot of normalized H2B intensity versus recovery time shows little correlation between chromatin structure and diffusion. (C) Correlation coefficients between recovery time and H2B intensity for N=11 nuclei show no significant correlation. (D) Peaks in recovery time for WT nuclei (n=9 cells, n=13 peaks) and TSA-treated nuclei (n=10 cell, n=12 peaks). No significant difference in recovery time is observed. Red lines represent mean and SEM.

While recovery times provide a useful means of comparing two conditions in a given experiment, determination of the diffusion coefficient is far more useful for comparison of results across experiments and techniques (4). Extraction of diffusion coefficients from FRAP experiments requires careful modeling of the bleached geometry; SPIM-FRAP is no exception. We then developed a simulation of diffusion during a SPIM-FRAP experiment (Fig. 3, Movie S2. See Supplementary Materials for a full derivation). The primary assumption in our model is that diffusion in and out of the image plane is the dominant source of the recovery of the fluorescence signal. This simplifies the full 3D problem to a 1D approximation governed by the equation

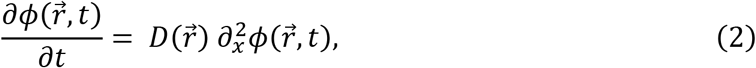

where 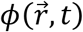 is the concentration of bright molecules and 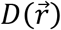 is the spatially-dependent diffusion coefficient. We used the theoretical profile of our Line Bessel Sheet to determine the region that will be photobleached. We additionally verified that we are bleaching a sheet consistent with this profile (Fig. S2, Movie S3). Furthermore, the common assumption that the photobleach pulse is significantly shorter than the relevant time scale does not hold for our case (*τ*_*D*_ ≈ 46 ms for 1D diffusion where *τ*_*D*_ = *l*^2^/2*D*, *l* = 675 nm is the full width at half maximum of the light sheet and *D* = 5 *μm*^2^/*s*), so it was necessary that we account for diffusion into the region being photobleached during the photobleach pulse. We performed this simulation across a sweep of diffusion coefficients, then fit Eq. 1 to the recovery of the simulation to determine the corresponding values of *B* and Τ for a given diffusion coefficient (Fig. 3C). The measured peak recovery times for WT cells range from 0.209 s to 0.832 s with a mean of 0.478 s. according to our simulation, this means we observed diffusion coefficients ranging from 2.22 μm^2^/s to 21.6 μm^2^/s with a mean of 4.52 μm^2^/s. The simulated values of recovery percentage are also consistent with our experiments. We can use these results to convert experimental maps of *τ* to maps of *D* (Fig. 3D). Previous literature on the intranuclear diffusion of small molecules gives diffusion coefficients ranging from approximately 4 μm^2^/s to 50 μm^2^/s (8–10, 18–23). Our results then fall on the lower end of the previous reported values. This could potentially be due to factors regarding simulation, such as the principle assumptions, a dependence of *D* on the geometry of the nucleus, or the specific bleach correction used in our analysis (4). Alternatively, the slight discrepancy could be due to the addition of the nuclear localization sequence to the GFP molecule which subsequently changes RNA binding and molecular weight (17).

**Figure 3.**
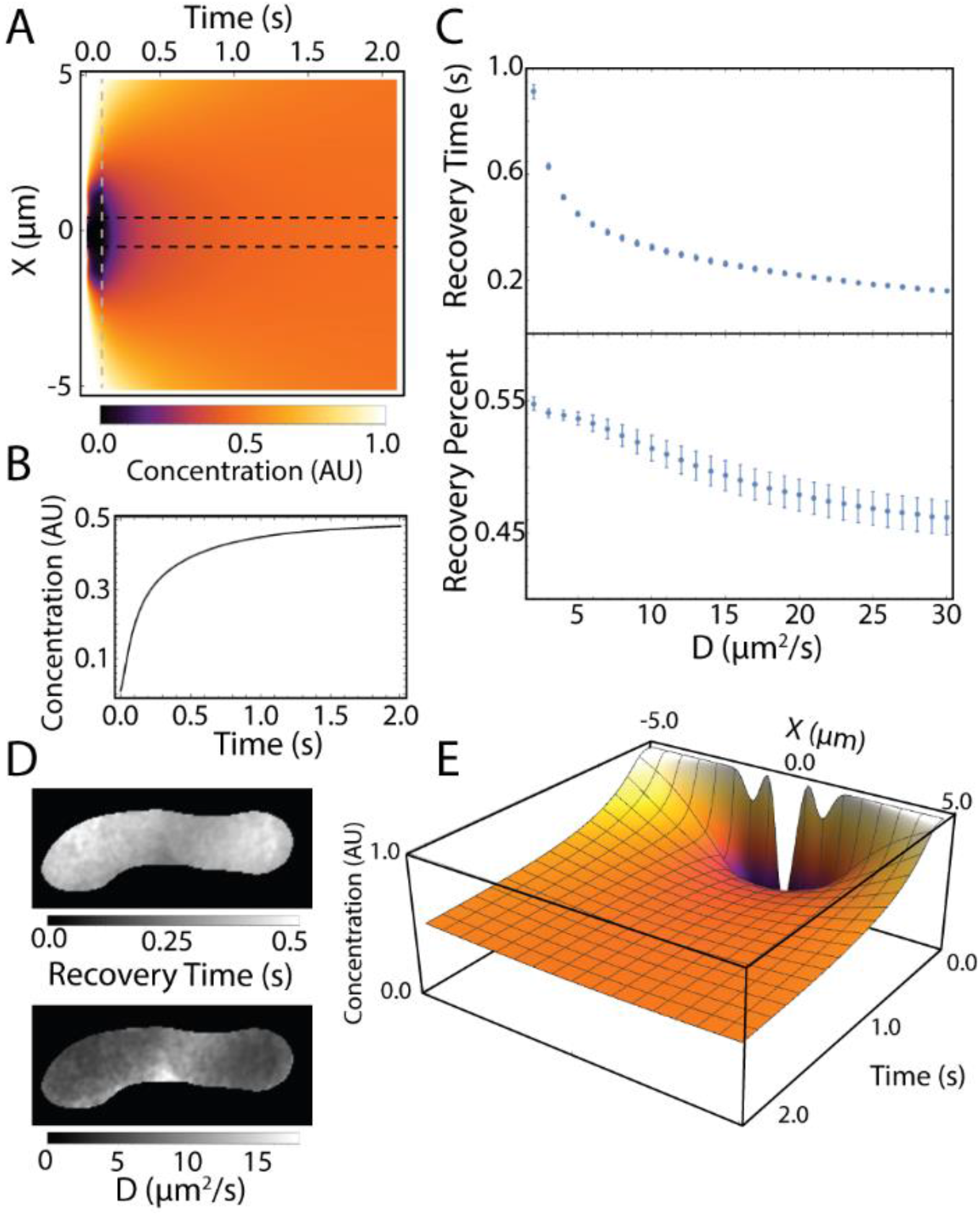
Diffusion simulation connects measured recovery times to diffusion coefficients. (A) Simulated diffusion for 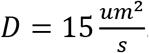. The grey dashed line represents when the light sheet turns off. Black dashed lines represent the region being integrated to determine recovery times. (B) Integrated concentration between the black dashed lines in (A) as a function of time (Movie S2). (C) Plot of recovery time versus diffusion coefficient for the simulated SPIM-FRAP diffusion process. Plot of recovery percentage versus diffusion coefficient for the simulated SPIM-FRAP diffusion process. Error bars represent standard errors for parameter fits. (D) An experimental map of recovery time for NLS-GFP and the corresponding map of *D*. (E) A 3D rendering of the diffusion simulation. At t=0, the initial concentration is the inverse of the light sheet profile.

## Conclusions

We have presented a unique combination of SPIM and FRAP that proves useful for making simultaneous FRAP measurements for each pixel in a given 2D plane. This allows one to studying heterogeneous diffusion and protein recovery on timescales ranging from milliseconds to hours. Such measurements were not previously accessible by traditional FRAP experiments and are an order of magnitude faster than FCS-based techniques. As with any development in methodology, SPIM-FRAP has limitations which are described in the supplemental material. We demonstrate that SPIM-FRAP can be used to track intranuclear diffusion of small molecules, specifically NLS-GFP. The recovery times show heterogeneity across the whole nucleus that is uncorrelated with histone density. Intranuclear diffusion also seems to be independent of chromatin compaction levels, pointing to another source of heterogeneous distribution such as intranuclear liquid-liquid phase separation, variable concentration of macromolecules, or binding of NLS to RNA. We have also shown through a 1D diffusion simulation that SPIM-FRAP produces diffusion coefficients that are consistent with previously reported values. SPIM-FRAP is poised to be immediately implemented on almost any light sheet microscope with minimal software development, making it a new tool for biologist to study not only the timescales and magnitudes of protein turnover and diffusion, but the spatial distributions as well.

## Author Contributions

C.M.H conceived of the SPIM-FRAP method and performed all experiments, simulations, and analyses. E.T.O performed the cell culture and sample preparation. M.R.F. and R.S. oversaw the project. All authors wrote the manuscript.

## Acknowledgements

We would like to thank the Lammerding Lab (Cornell University) for generously donating the cell lines used in this study and Talley Lambert for providing helpful feedback during the drafting of the manuscript. C.M.H. is supported by the NSF GRFP (DGE-1650116) and the Caroline H. and Thomas Royster Fellowship. E.T.O, M.R.F, and S.R are supported by NIH and NSF (NSF/NIGMS 1361375) as well as NIH (NIBIB P41-EB002025).

## Supporting Citations

References (24–27) appear in the Supporting Material.

## Supplementary Materials and Methods

### Cell Culture and Sample Preparation

Two MDA-MB-231 cell lines, transfected with NLS-GFP and either H2B-mCherry of 53BP1-mCherry, were a generous gift from the Lammerding Lab. Complete transfection protocols and reagents can be found in prior publication (1). Cells were cultured in DMEM/F12 with 10% FBS (Sigma-Aldrich) and 1X antibiotic antimycotic (Gibco) without phenol red. The media has 15 mM Hepes buffer which helps stabilize the pH during experiments. One day before the experiment, 50-70% confluent cultures were trypsinized and plated on polyacrylamide gels such that only 1-3 cells were present per field of view at 60x magnification. Polyacrylamide gels were used in order to eliminate reflections during side-view imaging. They were made with high stiffness (55 kPa) as described in our previous work (2) and coated with collagen as a final extracellular matrix protein. Briefly, 10 μL of activated gel solution was deposited on APTES-treated 40 mm round coverslips and a 22×22 mm square coverslip was quickly placed on top. The top coverslip had been treated with HMDS via vapor deposition to facilitate easy removal after polymerization. The gel included 1% polyacrylacrylic acid to provide carboxylic acid groups within the gel. This promoted adhesion to the APTES coated glass substrate, and reactive sites for attachment of collagen after gelation. After gelation and coverslip removal under deionized water, the gel was allowed to dry briefly such that a 10 mm diameter glass cloning cylinder (316610, Corning) lightly coated with vacuum grease (1597418, Dow Corning) could be secured. Immediately, a solution of 10 mg/mL EDAC and 1 mg/mL NHS in phosphate buffered saline (PBS) was placed into the cloning rings, and the assembly was placed into sterile plastic petri dishes. The dishes were then placed in a 37⁰C chamber at 100% humidity for 15 minutes. The EDAC buffer was then replaced twice with PBS at room temperature, and then with 50 ug/mL collagen (Rat Tail Type I, Invitrogen) for 30 minutes at 37⁰C. The collagen solution was then replaced with PBS twice, and then with DMEM F12 growth media. Samples were then placed in the cell culture incubator to equilibrate at least 30 minutes before cells were added. For treatment with Trichostatin-A (TSA), TSA was dissolved to 10 mM in DMSO, and then serially diluted in PBS to 4 μM on the day of treatment. 10 μL of the 4 μM solution in PBS was then added to the cells as they were growing in 190 μL of media in 10 mm cloning cylinders, for a final concentration of 200 nM. Experiments were carried out 24-28 hours after drug addition. The 2×10^(−5) dilution of DMSO, giving 0.002% v/v final concentration was judged to be insignificant to the TSA effect.

### Image Acquisition and Analysis

Live MDA-MB-231 cells co-expressing either NLS-GFP and H2B-mCherry or NLS-GFP and 53BP1-mCherry were plated on 55 kPa polyacrylamide gels one day prior to examination on our custom light sheet microscope (3). We utilized vertical light-sheet based illumination and side-view imaging via a reflective prism adjacent to a cell of interest to first collect side-view (Y-Z) light-sheet fluorescence images. For cells expressing H2B-mCherry, we first collected one image of the H2B at an exposure time of 200 ms. We then collected 100 images of the NLS with a 5 ms exposure time and 5 ms readout time. At this point, a single Y-Z sheet was bleached for 100 ms with high-intensity 488 nm light. Immediately after the vertical sheet was bleached, an additional 300 images of the NLS were collected at the same exposure and readout time. The laser power was measured to be 1.50 ± 0.03 mW for the bleach pulse and 53.3 ± 0.7 μW for the standard image acquisition.

Image sequences were loaded into FIJI (4), and the first image was used to generate a mask of the nucleus as described in our previous work (2). The image sequence was subsequently blurred using a 1-pixel Gaussian blur. All further analysis is performed in Wolfram Mathematica 11.2. The first 100 images were used to correct the time series for photobleaching via an exponential bleach correction. This photobleaching correction was applied to all images in the time series and was not performed on a pixel-by-pixel basis but rather uniformly across the whole image. There is potential for the photobleaching to be spatially dependent along the axial direction due to dispersion of the light sheet, however this effect is negligible in our work as our imaging conditions seek to minimize photobleaching and the depth of field of the light sheet is greater than the height of the nuclei. For each pixel in the mask of the nucleus, an exponential recovery curve of the form 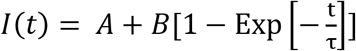 was fit to the first 200 intensity values of that pixel immediately after the bleaching step. From each curve, we extracted both the characteristic recovery time, *τ*, as well as the recovery percentage, 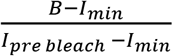.

Finally, we sought to verify the geometry of our photobleaching. To do so, we photobleached a single plane of a live MDA-MB-231 nucleus expressing H2B-mCherry. H2B has a recovery time on the order of hours for mammalian cells (5), so this sample was chosen such that we could observe the bleached region without concern for recovering too quickly. After photobleaching a single plane, we collected volumetric images with our custom SPIM microscope (Figure S2). We observed a clear photobleached plane through the center of the nucleus as well as slight photobleaching from the concentric side lobes characteristic of a Line Bessel Sheet. These side lobes are accounted for in our diffusion simulation. We did not observe any significant bleaching outside of the light sheet region due to scattered light. This demonstrated that photobleaching with a light sheet, and subsequently SPIM-FRAP, is indeed a reliable and reproducible technique.

### SPIM-FRAP Simulation

To validate our analysis protocol, we simulated SPIM-FRAP experiments based upon our experimental measurements (Figure S1). Simulated data sets were generated by first calculating the mean and standard deviation of both the recovery time and recovery percentage as well as the standard deviation of the plateaued recovery curve. For a given nucleus, the NLS image immediately after the bleach pulse was used as the starting image. Each pixel was prescribed an exponential recovery of the form 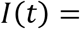 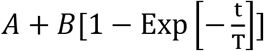 with a specified recovery time and recovery percentage based upon the mean of the experimental data. We simulated 3 conditions to isolate the contribution of each aspect of our analysis pipeline. First, we added noise to the simulated recovery based on the noise in the experimental data and fit an exponential recovery curve to each pixel. Next, we added a 1-pixel Gaussian blur along with the additional noise before fitting the recovery. Note that this serves to tighten the distribution of recovery times at the cost of spatial resolution. Finally, we added a predetermined spatial pattern of the recovery times in the form of 5×5 pixel squares with recovery times 20% lower than the rest of the pixels. We were able to clearly discern this structure after the addition of noise and Gaussian blurring. That is, the prescribed 20% change is well above the noise floor of our technique. These simulations can be compared to the experimental data set to validate the heterogeneity present in the map of recovery times.

### Diffusion Simulation Theory

Traditionally, FRAP experiments use simple bleaching geometries and minimize bleaching times in order to use analytical models to convert recovery time to a diffusion coefficient (6). Our light sheet microscope employs a Line Bessel Sheet (LBS), which features additional side lobes concentric to the main central lobe (3). This prevents us from applying any model with a simplified geometry. Furthermore, our bleach time of 100 ms is on the order of the measured recovery time; this means that we must account for diffusion occurring during the bleach pulse itself. To understand how our measured recovery times corresponded to diffusion coefficients, we computationally modeled diffusion in our system. The full three-dimensional diffusion equation is given by

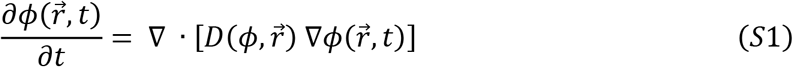

where 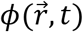 represents the concentration of bright molecules as a function of space and time, and 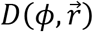 represents the diffusion coefficient as a function of concentration and space.

#### Assumption 1

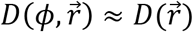. Justification: First, the concentration of NLS-GFP in the nucleus is effectively uniform, so there is little variation in *ϕ* which subsequently means there is little change in *D* due to variation in *ϕ*. Additionally, the concentration is that of dark versus bright molecules, whether or not the molecules are fluorescing has no physical bearing on the local diffusion coefficient and therefore 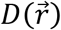 is effectively independent of *ϕ*. It is important to note that this assumption may not hold true for all experiments, and care should be taken in considering this assumption when implementing SPIM-FRAP for other studies.

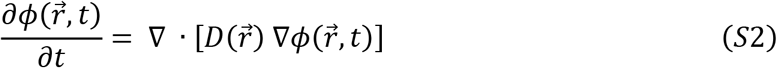

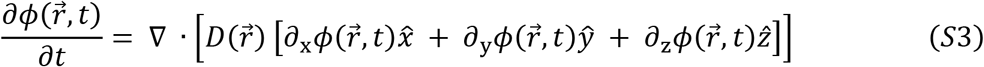

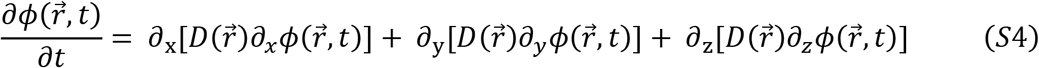

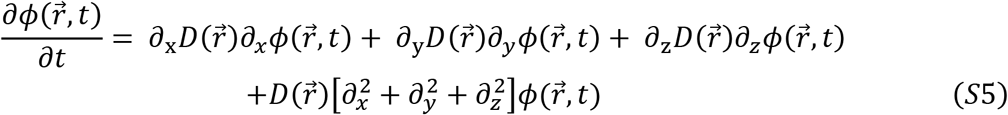

#### Assumption 2

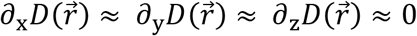 Justification: We observe change in the diffusion coefficient on the order of only a factor of two across the entire nucleus, meaning the local changes in the diffusion coefficient are negligible.

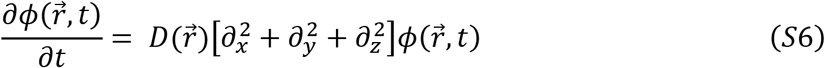

#### Assumption 3

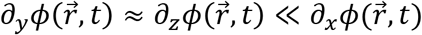. Justification: The primary plane of symmetry being broken is that of the x direction because the bleached pattern forms a y-z sheet. Diffusion in the y and z directions will then cause a far smaller change in concentration than diffusion in the x direction. This may not hold true for samples with larger spatial heterogeneity than NLS-GFP, which would subsequently require more detailed modeling.

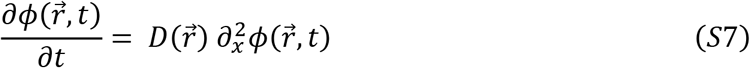

#### Initial Conditions

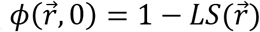 where 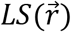 represents the normalized, theoretical profile of the light sheet.

#### Bleaching Conditions

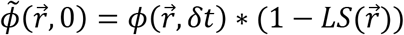. That is, Equation S7 is iteratively solved during the bleaching time and at each iteration the concentration was multiplied by the inverse profile of the light sheet. The new concentration profile was then used as the initial condition for the next iteration.

#### Boundary/Normalization Conditions

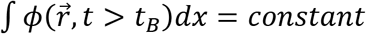. That is, once the bleaching has stopped, the total concentration of bright molecules remained constant.

### SPIM-FRAP Limitations

As with any advancement in methodology, there are accompanying limitations. One of the immediate drawbacks of SPIM-FRAP in its current implementation is in movement of the sample on the timescale of the recovery being measured. If the sample were to move into or out of plane, a false recovery would be detected (Figure S3). SPIM-FRAP with accompanying volumetric imaging can remedy this issue as one can monitor the bleached region even if it were to move in space; SPIM-FRAP with fixed plane imaging, however, is limited to measuring dynamics that are faster than cell morphodynamics and motility. Additionally, one must carefully consider the light sheet’s depth-of-field, defined to be the length scale in the direction of propagation for which the light sheet has minimal dispersion. If the depth-of-field is comparable the size of the sample, dispersion of the light could conflate the quantification. Here, we are implementing a light sheet with a theoretical depth-of-field >10 μm (7) while the height of the nuclei is generally ~5 μm. Hence, our recovery maps do not show a systematic trend in the direction of propagation of the light sheet. Our vertical light sheet system allows us to utilize shorter light sheets. Other geometries must be cognizant of this upon implementation of SPIM-FRAP. Light sheets are also subject to striping artifacts (Figure S4), which could further complicate measurements or make them infeasible. Finally, the presented work presumes that the concentration of bright fluorophores is effectively constant throughout the nucleus; this is not entirely the case. The distribution of NLS-GFP throughout the nuclear volume that is not constant, and this may have implications for our quantification. However, the variation of the distribution of NLS-GFP is far smaller than the variation induced by the photobleaching. This may not be true for all samples, and this should be considered in future experiments. Despite the aforementioned limitations of SPIM-FRAP, the benefits of resolving spatial heterogeneity with FRAP and the order of magnitude improvement of acquisition time relative to FCS prove useful for furthering the field of diffusion and protein dynamics.

## Supplemental Figures

**Figure S1.**
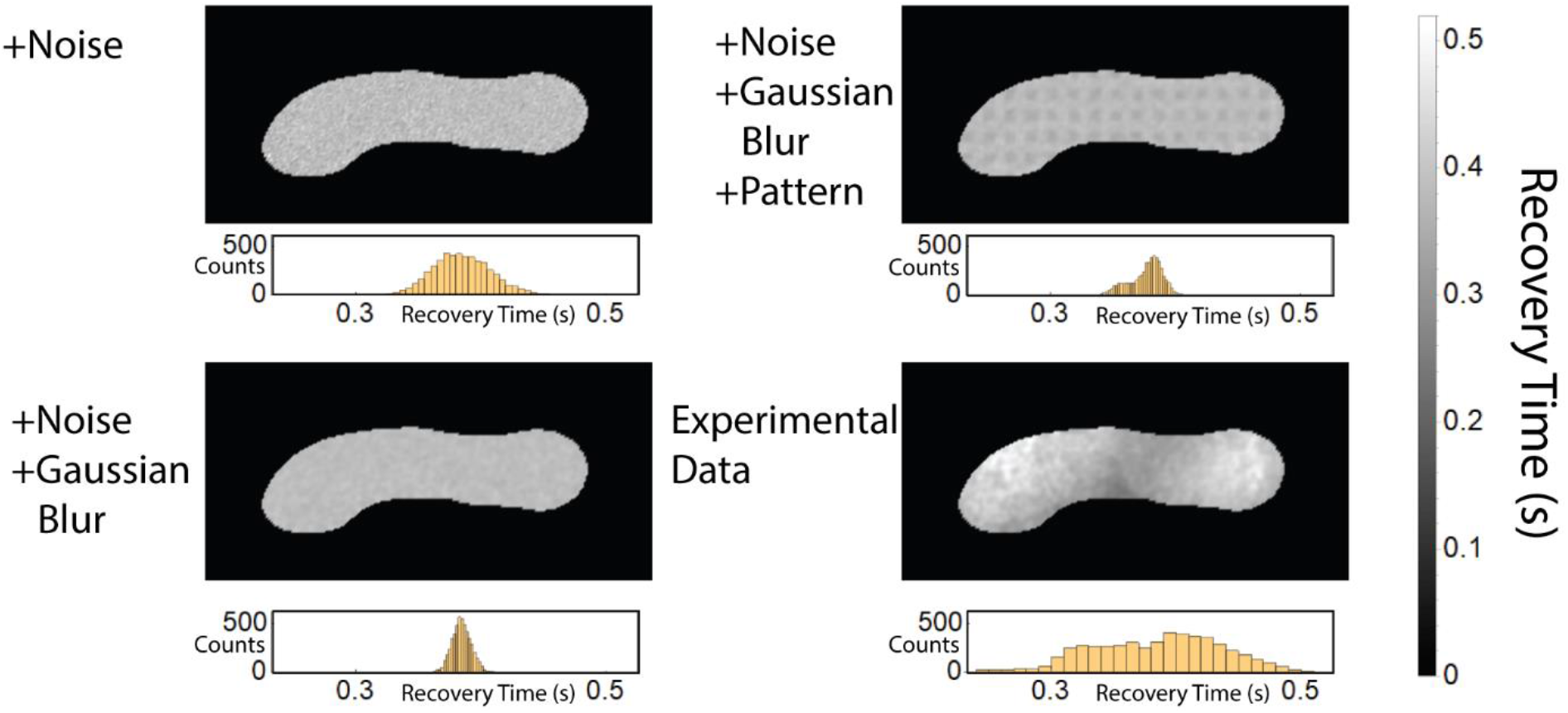
SPIM-FRAP simulated data sets. (Top, Left) Simulation with noise in the recovery curve. Ground truth is a uniform recovery time. (Bottom, Left) Simulation with noise in the recovery curve and Gaussian blurred before analysis. Ground truth is a uniform recovery time. (Top, Right) Simulation with noise in the recovery curve and Gaussian blurred before analysis. Ground truth is a 5×5 pixel array of recovery times that differ by 20%. (Bottom, Right) Experimental data set.

**Figure S2.**
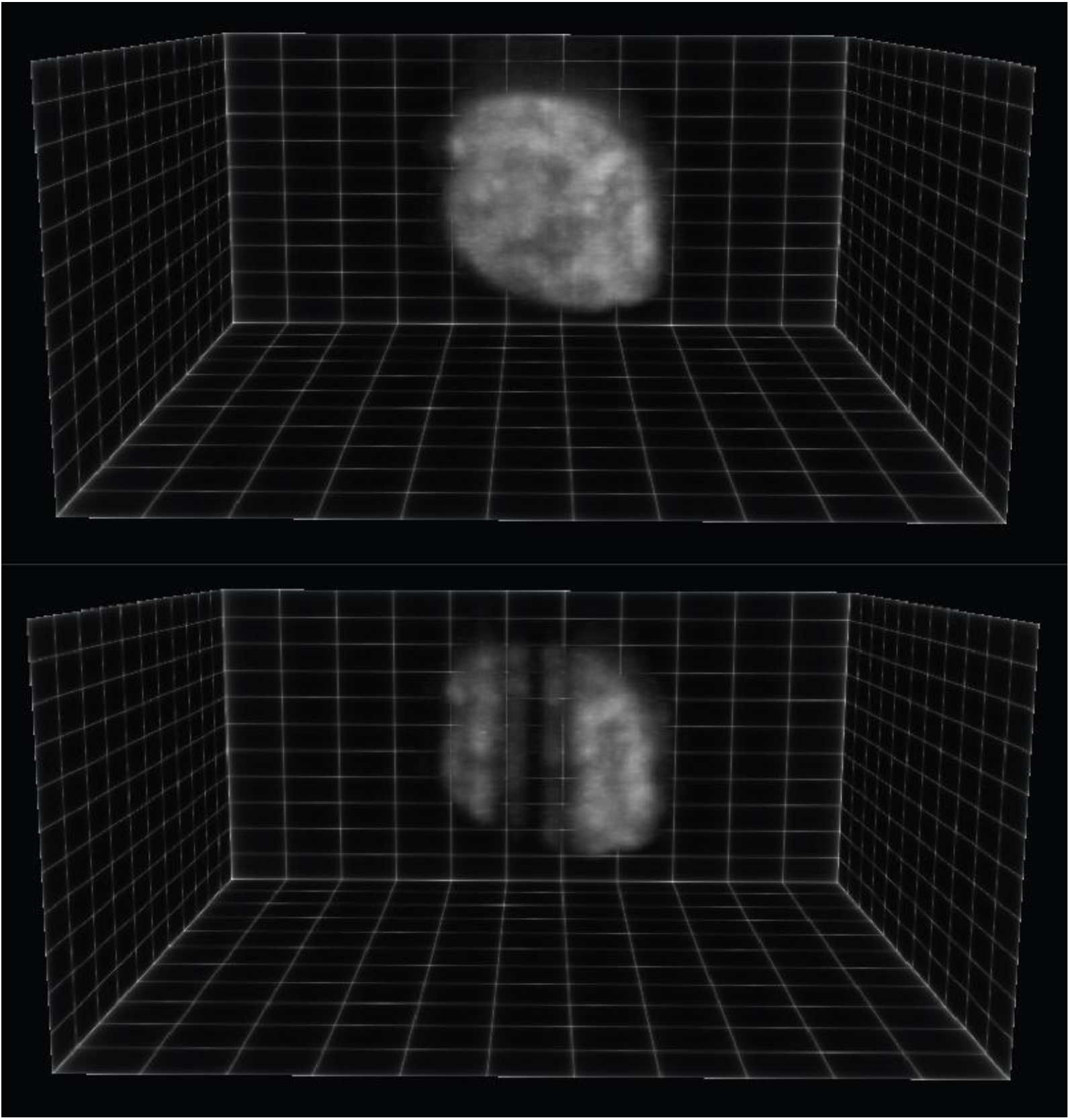
3D rendering of a live MDA-MB-231 nucleus expressing H2B-mCherry both pre and post SPIM photobleaching. A clear 2D plane is bleached through the nucleus as well as small photobleaching of the concentric side lobes. See Movie S3 for additional views.

**Figure S3.**
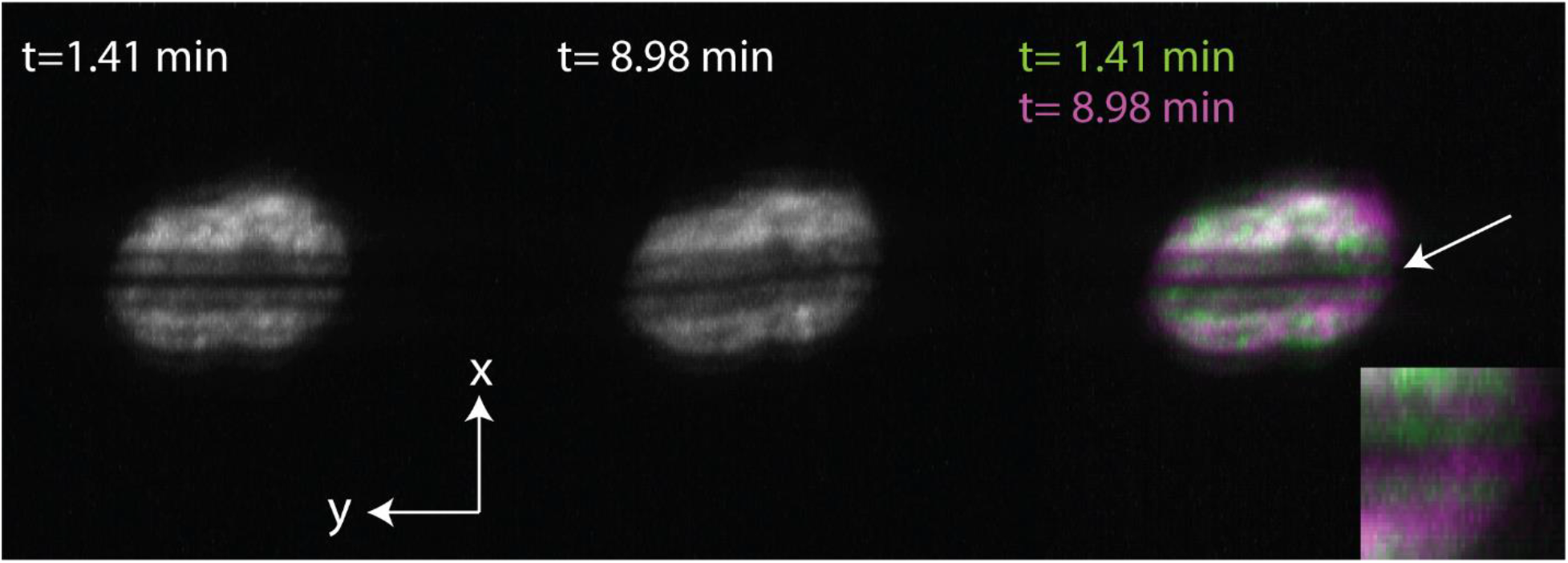
Maximum intensity projections of the reconstructed X-Y view of a live MDA-MB-231 nucleus expressing H2B-mCherry at different time points. The merged view shows the location of the bleached region moves in the x direction, which could conflate recovery measurements.

**Figure S4.**
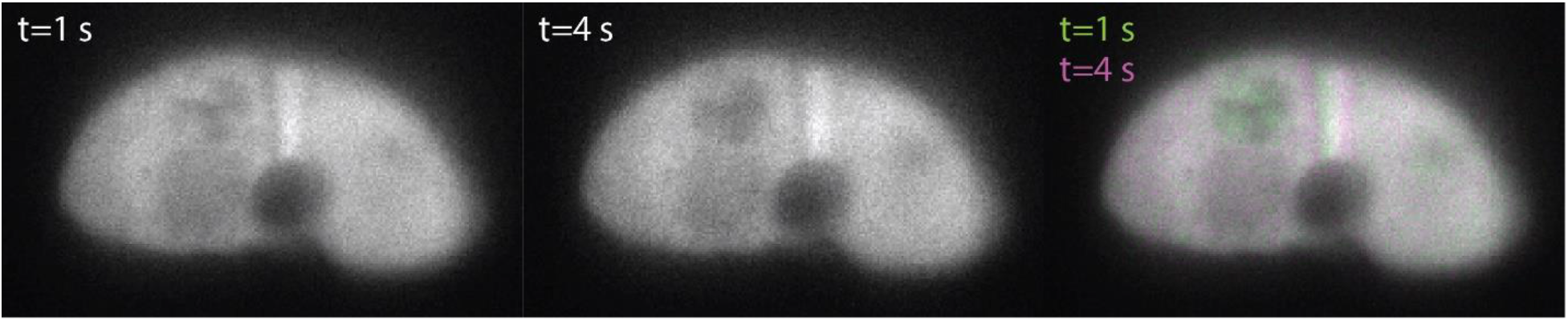
Selected images from an unsuccessful SPIM-FRAP experiment due to striping artifacts from the light sheet. The vertical stripe caused by the nucleolus shifts in time due to cell movement, making the recovery curves unusable.

## Supplementary Movie Captions

**Movie S1.** A side-view SPIM-FRAP image sequence of NLS-GFP showing the simultaneous recovery across the entire image plane.

**Movie S2.** A time series of the1D diffusion simulation, showing how concentration develops dynamically as a function of time.

**Movie S3.** 3D rendering of a live MDA-MB-231 nucleus expressing H2B-mCherry both pre and post SPIM photobleaching. A clear 2D plane is bleached through the nucleus as well as small photobleaching of the concentric side lobes. See Movie S3 for additional views.

